# NMR assignments and secondary structure analysis of the human 5MP1 C-terminal domain

**DOI:** 10.64898/2026.08.11.744028

**Authors:** Ayşenur Şeker, Simran Anand, Assen Marintchev

## Abstract

Eukaryotic translation initiation is tightly regulated by interactions among translation initiation factors (eIFs) that ensure accurate start codon selection. The translation regulator, eIF5 mimic protein 1 (5MP1) contributes to this process by competing with eIF5 for binding to eIF2, thereby increasing the stringency of translation initiation. Despite its important regulatory role and emerging involvement in tumorigenesis, structural information on human 5MP1 remains limited. Here, we report the near-complete backbone and partial side-chain NMR resonance assignments of the C-terminal domain of human 5MP1 (residues 250–419), carrying a W404E substitution that disrupts dimerization. The WT protein forms a dimer at NMR concentrations, which increases the effective size of the protein and also causes disappearance of peaks corresponding to aminoacids at the dimer interface due to conformational exchange.

Backbone resonance assignments were completed for 96.4% of the non-proline residues. Secondary structure was analyzed using Chemical Shift Index (CSI) and compared with the AlphaFold structural model. Regions of disagreement between the experimental and computational secondary structure assignments were further examined using ^15^N-NOESY-HSQC spectra, allowing experimental validation of local structural features. While the AlphaFold model accurately reproduces the overall fold of the 5MP1 C-terminal domain, several localized discrepancies were identified, particularly near the N- and C-terminal regions of the domain, where experimental NMR data support alternative secondary structure assignments. These resonance assignments and experimentally validated structural features provide a foundation for future investigations of the molecular interactions, dynamics, and functions of 5MP1 in translation initiation.

## Biological Context

Eukaryotic translation initiation is a strictly controlled, multi-step process involving a network of proteins called eukaryotic translation initiation factors (eIFs). Interactions among these proteins and with the small ribosomal subunit determine the efficiency and fidelity of translation. The eIF5-mimic protein 1 (5MP1) is an important component of this network, playing a role in the stringency of translation initiation by competing with eIF5. eIF5 is the GTPase-activating protein for eIF2, the factor that delivers the initiator Met-tRNA_i_ to the translation pre-initiation complex (PIC), which scans the mRNA for a start codon within an appropriate sequence context (reviewed in (Marintchev, 2025).

5MP1 was formerly known as Basic leucine zipper and W2 domain 2 (BZW2). However, later it was predicted by us (Marintchev & Wagner, 2005) and AlphaFold (Varadi et al., 2022) that the N-terminal domain of 5MP1 is an MA3 HEAT domain homologous to the second HEAT domain of eIF4G, not a leucine zipper. Its C-terminal domain is a W2 HEAT domain homologous to eIF5-CTD (Koonin, 1995, Singh et al., 2011). The competition between 5MP1 and eIF5-CTD for binding to eIF2 results in increased start codon selection stringency. This dynamic interplay modulates the repression of non-AUG initiation (Tang et al., 2017). Thus, 5MP1 autoregulates its own expression and is also cross-regulated with eIF5 (Loughran et al., 2018).

5MP1 has been reported to be involved in tumorigenesis in various cancer types (Kozel et al., 2016, Li et al., 2025). Upregulation of 5MP1 is associated with modulating tumor cell growth, making it a potential prognostic biomarker and therapeutic target (Li et al., 2021). However, the limited understanding of its interactions and structural properties has impeded further research.

Here, we used NMR spectroscopy to obtain the NMR backbone resonance assignments of the 5MP1 C-terminal domain (CTD) using a set of standard triple-resonance experiments (reviewed in (Marintchev et al., 2007)). The wild-type protein dimerizes at concentrations required for NMR, forming a ∼40 kDa complex, which also causes disappearance of peaks corresponding to aminoacids at the dimer interface due to conformational exchange. Therefore, the monomeric W404E mutant was used for this study.

## Methods and Experiments

### Protein Expression and Purification

For recombinant protein production, a codon-optimized DNA sequence of human 5MP1-CTD-W404E, corresponding to residues 249-419 carrying the W404E mutation, was inserted into a modified pET21a (Novagen) expression vector. This construct was engineered to express the target protein with an N-terminal tag composed of a GB1 domain, a hexahistidine (His_6_) sequence, and a Tobacco Etch Virus (TEV) protease cleavage site.

The protein was expressed in *Escherichia coli* BL21(DE3) cells. For isotopic labeling, cells were grown in 1 L of minimal medium containing ^15^NH_4_Cl and ^13^C-glucose as the sole nitrogen and carbon sources, respectively. These cultures were grown at 37°C until reaching an OD_600_ of 0.6–0.8. Protein expression was then induced with 1 mM IPTG overnight at 20°C.

Cell pellets from 1 L cultures were resuspended in 40 mL of lysis buffer (10 mM sodium phosphate pH 7.0, 300 mM KCl, 7 mM β-mercaptoethanol, 0.01% NaN3) supplemented with 0.1 mM AEBSF, a protease inhibitor cocktail, and 1 mg/mL lysozyme. Lysis was performed by sonication, and the resulting lysate was clarified by centrifugation to remove insoluble debris.

The soluble, tagged protein was isolated from the supernatant via pre-equilibrated Talon CellThru affinity resin at 4°C. The resin was washed thoroughly with a running buffer (300 mM KCl, 10 mM sodium phosphate pH 7.0, 7 mM β-mercaptoethanol, 0.1 mM AEBSF, 0.01% NaN3) and subsequently with the same buffer containing 4 mM imidazole. The bound protein was eluted with running buffer containing 200 mM imidazole, and the collected fractions were immediately supplemented with 2 mM DTT and 1 mM EDTA.

The eluted protein was further purified by ion-exchange chromatography using a 5 mL HiTrap Q FF column. The column was equilibrated in a low-salt buffer (100 mM NaCl, 10 mM sodium phosphate pH 7.0, 1 mM DTT, 1 mM EDTA, 0.1 mM AEBSF, 0.01% NaN_3_), and the protein was eluted using a 25 mL linear gradient to 50% of a high-salt buffer (1 M NaCl). To remove the N-terminal tag, the purified protein was incubated with TEV protease at a 1:50 mass ratio for two hours at room temperature in a buffer of 150 mM NaCl, 10 mM sodium phosphate pH 7.0, 1 mM DTT, and 1 mM EDTA. This cleavage leaves a single N-terminal glycine residue from the tag. A final ion-exchange step, using the same protocol as before, was performed to separate the cleaved tag and TEV protease from the final product.

### NMR Spectroscopy

All NMR experiments were conducted on a Bruker AMX 500 MHz spectrometer equipped with a cryoprobe. For data collection, the purified protein was exchanged into an NMR buffer composed of 10 mM sodium phosphate (pH 7.0), 150 mM KCl, 2 mM DTT, 1 mM EDTA, and 5% D_2_O.

Backbone resonance assignments were achieved using a standard suite of triple-resonance experiments, including HNCO, HN(CA)CO, CBCAHN, CBCA(CO)NH, (H)C(CO)NH, HNCA and HN(CO)CA. Data was acquired at 25 °C.

All raw NMR data were processed using the NMRPipe software package (Delaglio et al., 1995). The subsequent visualization of spectra and completion of the backbone assignments were performed using CARA (Keller, 2004).

### Extent of Assignments and Data Deposition

The ^1^H–^5^N HSQC spectrum for the human 5MP1-CTD-W404E is presented in **Figure 1**, displaying well-resolved peaks characteristic of a folded protein domain. Several resonances were either missing or weak, which is typically attributed to residues undergoing conformational exchange or rapid proton exchange with water. The backbone resonance assignments achieved near-complete coverage. Specifically, 161 of the 167 non-proline amide correlations were successfully assigned, corresponding to an assignment completeness of 96.4%. Six non-proline residues could not be assigned: Gln 250, Glu 311, Glu 312, Lys 345, Val 346, and Ile 354. High completeness was also observed for other backbone nuclei: Cα resonances were assigned for 164 out of 170 residues (96.5%), and Cβ resonances were assigned for 157 of the 165 expected residues (95.2%). Carbonyl (CO) carbons were assigned for 162 out of 170 expected residues, reaching 95.3% completeness. Regarding side-chain assignments, 108 residues were expected to have assignments beyond the Cβ position, excluding aromatic rings, methionine methyl carbons, and arginine Cζ carbons. Within these 108 residues, a total of 187 side-chain assignments were anticipated, of which 82 were completed. This resulted in 44 residues having fully complete side-chain assignments and 22 residues having partial side-chain assignments.

**Figure 1.**
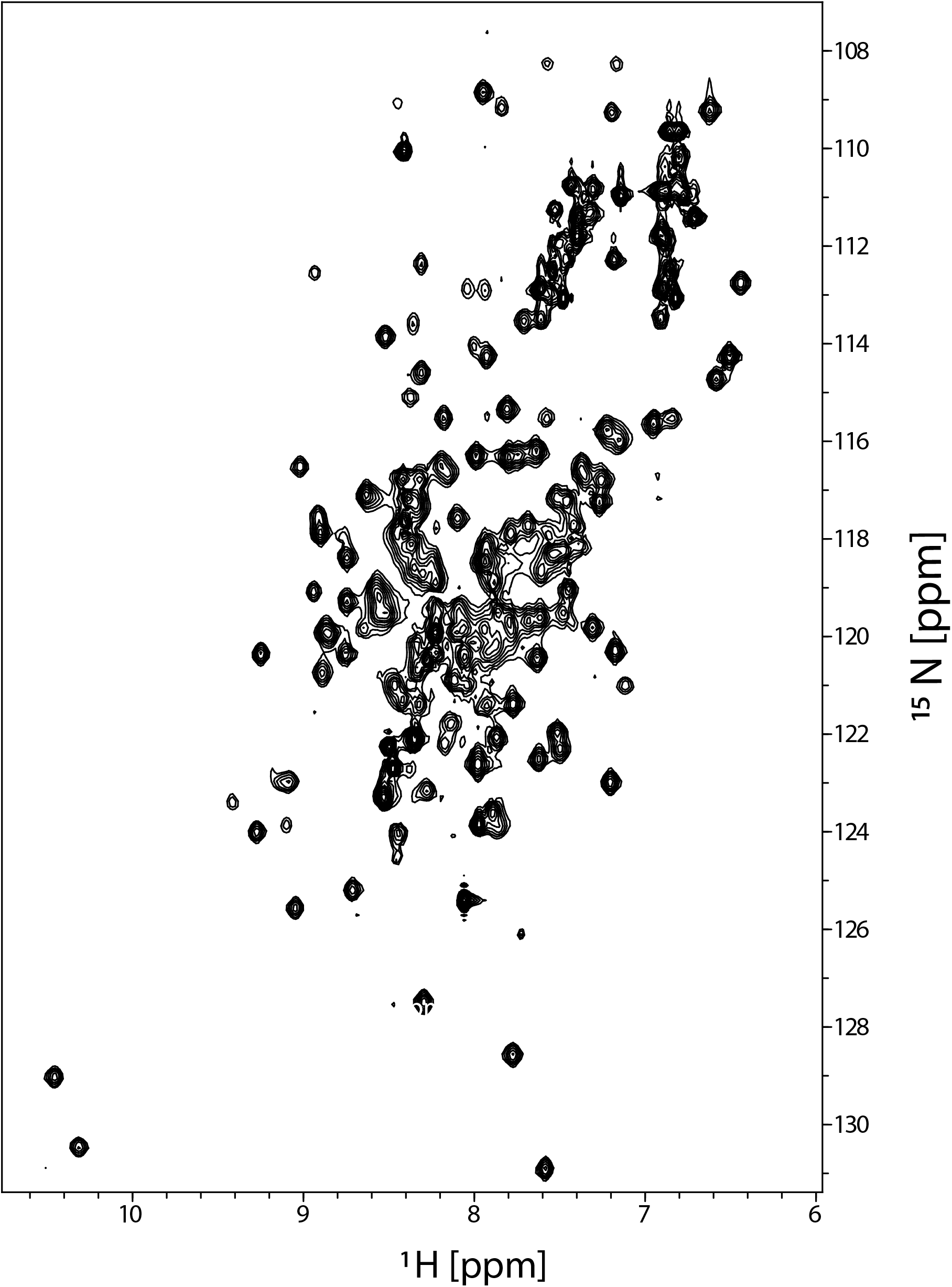
^1^H, ^5^N-HSQC spectrum for the human 5MP1-CTD-W404E fragment (residues 250– 419). Spectra were recorded at 25 °C at a field strength of 500 MHz, in a buffer containing 10 mM sodium phosphate (pH 7.0), 150 mM KCl, 2 mM DTT, 1 mM EDTA, and 5% D_2_O.

The comprehensive set of ^1^H, ^13^C, and ^15^N NMR chemical shift assignments for the human 5MP1-CTD-W404E fragment (aa. 250-419) has been deposited in the Biological Magnetic Resonance Data Bank (BMRB) under accession code 53383.

### Secondary Structure Analysis

Secondary structure was analyzed by integrating experimental NMR data with computational structural predictions made by AlphaFold. The backbone secondary structure was first evaluated using Chemical Shift Index (CSI) analysis with the CSI 3.0 server (Hafsa et al., 2015). CSI identifies secondary structure propensities by comparing experimentally determined backbone chemical shifts with residue-specific random coil reference values. For this analysis, the backbone chemical shifts of ^1^Hα, ^13^Cα, ^13^Cβ, and ^13^CO nuclei were used.

Based on these comparisons, residues are assigned scores corresponding to α-helical (+1), β-strand (−1), or random coil (0) conformations. Consecutive stretches of positive or negative values indicate regions with α-helical or β-strand propensity, providing an experimental assessment of local secondary structure.

The experimentally derived CSI assignments were subsequently compared with the AlphaFold structural model of human 5MP1. AlphaFold is a deep learning-based protein structure prediction method that predicts three-dimensional protein structures using the structures of homologous proteins as a template, or directly from amino acid sequences if structures of homologs are not available (Jumper et al., 2021, Varadi et al., 2022). Although AlphaFold has substantially advanced structural biology, computational predictions require experimental validation, particularly in regions that exhibit conformational flexibility or marginal secondary structure propensity.

Overall, the CSI analysis was in agreement with the AlphaFold model across most of the 5MP1-CTD. However, several localized regions displayed discrepancies between the experimentally derived secondary structure and the computational prediction.

To further evaluate these regions, three-dimensional ^5^N-edited NOESY-HSQC spectra were analyzed. Expected NH-NH cross-peaks were identified from the AlphaFold model based on predicted interproton distances. Prior to analysis of the regions of disagreement, reference α- helices showing agreement between the CSI analysis and the AlphaFold prediction were examined to calibrate the relationship between NH-NH distance and NOESY cross-peak intensity. Based on this calibration, NH-NH distances of ≤4.5 Å were considered sufficiently short to consistently produce observable NOESY cross-peaks under the experimental conditions.

Predicted NH-NH distances within regions where the CSI analysis and AlphaFold model differed were subsequently compared with experimentally observed NOESY spectra. Only reciprocal cross-peaks observed in both corresponding strips were considered supportive of the predicted structural model. Spectral overlap, weak diagonal resonances, or ambiguous peak assignments were considered insufficient for reliable interpretation.

Based on the combined experimental and computational analyses, regions of disagreement between the CSI assignments and the AlphaFold model were classified into three categories. Where the experimentally observed NOESY cross-peaks were consistent with the CSI analysis, the CSI/NOESY-predicted secondary structure was selected as the correct secondary structure (top secondary structure row in **Figure 2**, highlighted green). These comprised residues 252-257, the segment surrounding 294–295, 303, 337, and the C-terminal region 407–414. Since two independent experimental approaches support the same structural interpretation, these regions provide strong evidence that the local solution structure differs from the AlphaFold prediction. Where the experimentally observed NOESY cross-peaks were consistent with the AlphaFold model (bottom secondary structure row in **Figure 2**, highlighted yellow), the AlphaFold/NOESY-predicted secondary structure was selected (top secondary structure row in **Figure 2**). These included residues 290–293, 296–297, 326, 332, 382–383, and 391 - mostly at ends of helices and in segments with irregular secondary structure. Where insufficient or ambiguous NOESY information prevented further evaluation, the CSI-predicted secondary structure was selected as the correct secondary structure (top secondary structure row in **Figure 2**, highlighted grey). These regions included residues 286, 327–328, 336, the segment spanning 350–356, and 359.

**Figure 2.**
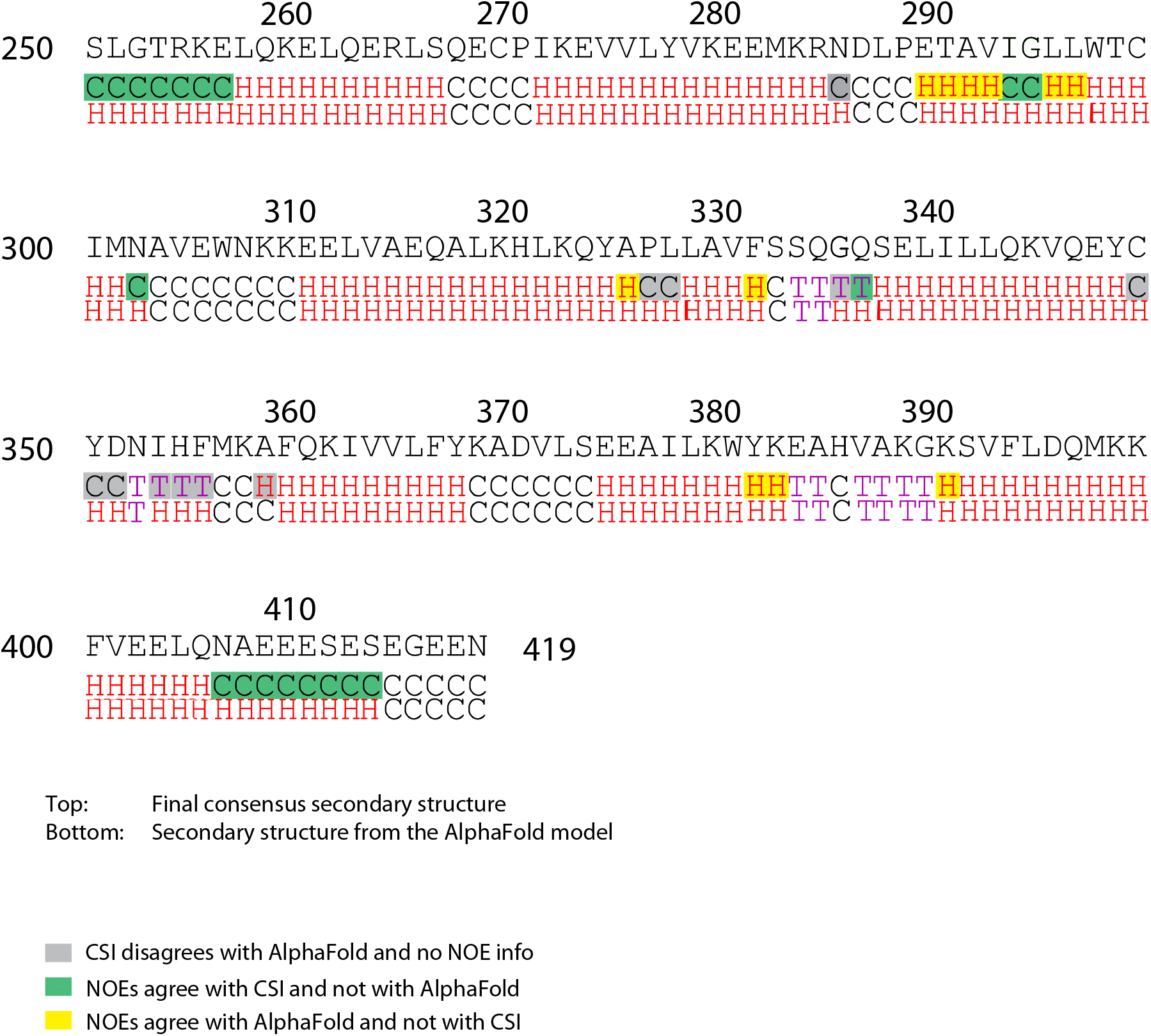
Comparison of the experimentally determined secondary structure and the AlphaFold structural model of human 5MP1-CTD-W404E (residues 250–419). The top row shows the final consensus secondary structure obtained by integrating the CSI analysis with the AlphaFold model and experimental NH–NH NOESY validation. The bottom row shows the original AlphaFold secondary structure prediction for comparison. Regions where NOESY data supported the CSI assignment over the AlphaFold prediction are highlighted in green. Regions where NOESY data supported the AlphaFold prediction are highlighted in yellow. Regions for which NOESY data were insufficient or ambiguous to distinguish between the CSI and AlphaFold assignments are highlighted in grey; in these cases, the CSI assignment was retained in the final consensus secondary structure.

## Conclusion

Here, we report the backbone and partial side-chain NMR chemical shift assignments for the C-terminal domain of the human 5MP1 W404E mutant (residues 250–419). The high completeness of these assignments provides a residue-specific framework for future structural and functional studies of 5MP1 and its interactions with components of the eukaryotic translation initiation machinery.

Chemical Shift Index (CSI) analysis of the experimental data confirmed the predominantly α-helical architecture expected for a W2 HEAT domain. To further evaluate the predicted structure, the experimentally derived secondary structure was compared with the AlphaFold structural model and independently assessed using NH–NH NOESY cross-peaks. Taken together, these analyses demonstrate that the AlphaFold model for the most part accurately reproduces the overall fold of the 5MP1-CTD, except for regions near the N- and C-termini of the domain, where it appears biased by the available structures of homologous proteins, and in the case of the C-terminus, possibly by crystal packing artefacts in the structure used as template. Importantly, the combination of CSI and NOESY analyses enables experimental validation and correction of the model.

Overall, the resonance assignments and structural analyses presented here establish a comprehensive experimental characterization of the human 5MP1 C-terminal domain. These data provide a foundation for future investigations of the molecular interactions of 5MP1 with translation initiation factors and illustrate the complementary roles of NMR spectroscopy and computational structure prediction in high-resolution protein structure determination.

## Statements and Declarations

## Acknowledgements

This work was supported by the National Institute of Health [GM134113 to A.M.]. We thank the NMR Core Facility at Boston University Chobanian & Avedisian School of Medicine for access to instrumentation and technical support.

## Author Contributions

A.S. performed NMR data analysis, assigned resonances, and conducted secondary structure analysis. S.A. performed NMR data analysis. A.M. collected NMR spectra, guided experimental design and supervised all stages of the project. A.S, S.A. and A.M. jointly interpreted the data and wrote the manuscript. All authors reviewed and approved the final version.

## Data Availability

The chemical shift assignments have been deposited in the Biological Magnetic Resonance Data Bank (BMRB) under accession number **53383**.

## Ethical Approval

The authors declare that no human or animal subjects were involved, and all experiments were conducted in compliance with institutional and ethical guidelines.

